# Ultrasonic Detection of Venous Gas Emboli and Tissue Gas in Rodents: Toward Imaging Biomarkers for Decompression Stress

**DOI:** 10.64898/2026.10.26.761998

**Authors:** Joshua B Currens, Katherine M Eltz, Rani S Sellers, Emmanuel Dugrenot, Virginie Papadopoulou

**Affiliations:** Department of Radiology, The University of North Carolina at Chapel Hill, NC, USA; Lampe Joint Department of Biomedical Engineering, The University of North Carolina at Chapel Hill and North Carolina State University, NC, USA; Department of Pathology Laboratory Medicine, The University of North Carolina at Chapel Hill, NC, USA; Divers Alert Network, Durham, NC, USA

## Abstract

**Introduction:** In humans, decompression sickness (DCS) risk assessment often relies on surrogate ultrasound detection of venous gas emboli (VGE). Rodents are widely used to study DCS and mortality, yet no established relationship exists between imaging and decompression outcomes. We exposed rats to dives with differing DCS risk and investigated whether ultrasound measurements reflected modeled DCS risk and revealed tissue-level manifestations of decompression stress.

**Methods:** Rats underwent dry saturation exposures to 700 KPA in a hyperbaric chamber while breathing either heliox (low-risk; pDCS = 0.1%) or air (high-risk; pDCS = 13%), where pDCS was estimated using an existing rodent DCS risk model. Following decompression, animals were anesthetized for serial cardiac and renal ultrasound imaging. Echocardiograms were evaluated for VGE using an ordinal scale. The mean grayscale intensity (arbitrary units, a.u.) was also quantified in renal region of interest (ROI). In a separate set of animals, histopathology was conducted to evaluate renal manifestations.

**Results:** Observed peak VGE grades increased with estimated DCS risk, from grade 1 at pDCS 0.1% to grade 2 at pDCS 13%. Multi-organ analysis found lower renal ROI echogenicity during periods of high VGE, with mean grayscale intensities of 67.0 a.u. and 60.7 a.u. for VGE grades 0-1 and 2-4, respectively. Renal histopathology identified round voids consistent with pre-mortem gas emboli.

**Conclusions:** VGE burden tracked with pDCS risk. The decrease in renal image intensity during high VGE may indicate ultrasound attenuation from acoustic shadowing caused by excess tissue gas bubbles. These findings advance the development of physiologically relevant imaging biomarkers of decompression stress.

## Introduction

Decompression sickness (DCS) is an inherent risk in diving operations and is thought to be caused by gas bubble formation in the tissues of divers, with clinical manifestations ranging from musculoskeletal pain and skin rash to paralysis or death.(1) During a dive, breathing gas is dispensed at the elevated ambient pressure, causing dissolved inert gases to equilibrate in the tissues. Upon ascent, the dissolved inert gas can form bubbles that enter venous circulation, which are typically filtered at the pulmonary capillary bed.(1,2) Circulating decompression bubbles, or venous gas emboli (VGE), are commonly detected in ultrasound images of the right heart or other venous blood vessels after experimental dives. In human diving studies, VGE are widely used as a surrogate measure of decompression stress, as they are frequently detected even in the absence of clinical DCS.(3) Importantly, the absence of VGE is a strong indicator of decompression safety, whereas even high VGE grades have low positive predictive value for DCS(3,4). This relationship remains poorly characterized in rodent models.(5)

Since DCS is rare in humans and ethical considerations limit experimental exposure to high-risk dive profiles,(3,6–8) preclinical rodent models are commonly used in DCS research to study dive outcomes and test hypotheses prior to translation in humans.(9–13) A recent scoping review discussed the utility of ultrasound for decompression monitoring in rodent models.(5) However, the review identified several gaps in the literature, including inconsistent reporting of methods and a lack of example ultrasound images with VGE in rodents.(5) Establishing an experimental workflow for decompression trials with ultrasound monitoring could improve study reproducibility and provide the groundwork for mechanistic evaluation of decompression physiology with ultrasound imaging. Other ultrasonic methods have been utilized for gas presence analysis in previous studies, expanding beyond VGE detection into other organs. In 1984, Daniels *et al.* demonstrated the potential utility of ultrasound signal analysis from soft tissue, identifying potential associations with DCS in animals and humans.(14) Later, L’Abbate *et al.* measured the liver of rats and human divers with a clinical ultrasound system after hyperbaric exposure, finding that signal intensity increased over time post-dive, possibly due to gas bubble presence.(15) Separately, Swan *et al.* identified a dual frequency signal change in pig muscle during decompression that was consistent with *in vitro* microbubble measurements.(16) Currens *et al.* assessed porcine lymph node region brightness and identified a decreased region brightness associated with severe DCS cases, likely due to acoustic shadowing from excess gas accumulation in the lymph node and surrounding region.(17) Most recently, Eltz *et al.* found that increasing gas burden in cardiac and liver regions of bycaught sea turtles similarly resulted in darker regions overall.(18) Taken together, these studies indicate that quantitative ultrasound analysis may improve decompression stress estimates.

The utility of a wearable ultrasound monitor for divers has been proposed, which may allow for decompression bubble detection during a dive exposure.(19) Future development of this technology relies on an understanding of the connection between systemic gas presence and DCS risk, but the lack of established and standardized small animal model imaging methods inhibits progress towards this end. Therefore, we have designed a pipeline for rodent decompression trials with ultrasound measurements. A trial was conducted following two different dive exposures to evaluate the workflow. We implemented the Lillo & Parker(20) probabilistic DCS model for risk estimation, which has been previously cited for rodent study design.(5,20) Lastly, we expanded beyond echocardiogram measurements into the renal system by conducting a pilot image analysis. To our knowledge, this is the first use and evaluation of multi-organ ultrasound imaging in a rodent decompression model, incorporating cardiac VGE assessment adapted from the human gold standard and comparison across dive profiles of differing modeled DCS risk. Together, these methods establish a framework for investigating the relationship between circulating VGE and tissue-level manifestations of decompression stress, supporting the development of advanced ultrasound imaging biomarkers.

## Methods

### Ethics Statement

All animal experimentation was conducted with prior approval by the University of North Carolina’s Institutional Animal Care and Use Committee (Protocol ID: 23-168).

### Hyperbaric Exposures

Eight Sprague-Dawley rats (12 - 13 weeks old, 4 male and 4 female) were purchased from Charles River Labs. The animals were housed in a temperature-controlled room (22 ± 2°C) on a 12-hour light/dark cycle with food and autoclaved water provided *ad libitum*. Rats acclimated to the facility for one week prior to the start of experimental procedures (13 - 14 weeks old, average weight: 305.3±45g). Rodent hyperbaric exposures were conducted one at a time in a small animal hyperbaric chamber to simulate a dive (Fig. 1). The animal was awake and secured in an enclosure for the duration of the exposure. An acrylic porthole allowed for visual monitoring of the animal throughout the hyperbaric exposure. The chamber was connected to a high-pressure gas cylinder and manually pressurized at a rate of 202 kPa per minute and decompressed at 50 kPa per minute. All exposures were conducted to a maximum pressure of 700 kPa for a duration of 90 minutes from dive start until the beginning of decompression. Animals were randomly assigned evenly to one of two breathing gases: Air (79% N2, 21% O2) or Heliox (79% He, 21% O2). The choice of breathing gas served as a straightforward means of modifying the anticipated decompression stress during the simulated dive. A brief chamber flushing procedure was completed prior to the beginning of each dive profile by increasing chamber pressure to 405 kPa then decreasing to 304 kPa, repeated twice. The pressure profile was monitored using a dive computer (Teric, Shearwater Inc., Richmond, BC, Canada) placed in a container of water within the chamber.

**Figure 1.**
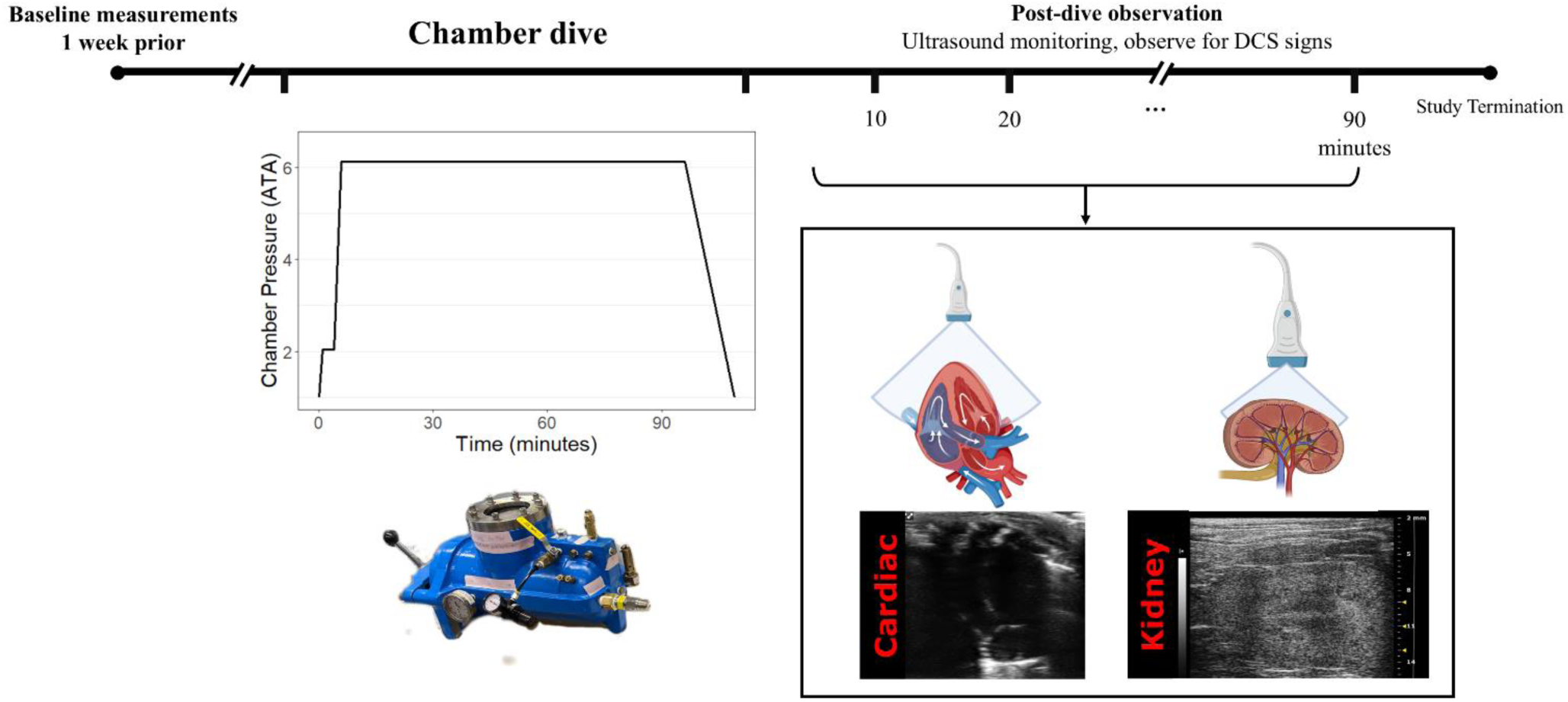
Decompression experimentation workflow. (a) Diagram of animal in hyperbaric with pressure profile below. (b) Example ultrasound imaging set up and animal position. Heart and kidney ultrasound cartoons in this image were created with BioRender.com.

### Post-Dive Observation

Upon returning to atmospheric pressure, the animal was immediately removed from the chamber and anesthetized with 5% isoflurane delivered in the assigned breathing gas at a flow rate of 1 liter per minute (L/min). Anesthesia was lowered and maintained at 2-2.5% isoflurane for the duration of the observation period, with adequate anesthetic depth confirmed by the absence of a toe-pinch reflex. The anesthetized animal was placed on a heating pad underneath a warming lamp to prevent hypothermia. Body temperature was monitored with a rectal thermometer. Animals were maintained under anesthesia for longitudinal ultrasound imaging for up to 90 minutes, after which they were euthanized according to the approved terminal study protocol. Animals were continuously monitored, and the study was terminated early if predefined humane endpoints were reached, including irregular or labored breathing, gasping or other signs of distress.

### Probabilistic Risk of DCS Estimation

The probabilistic DCS (pDCS) model developed by Lillo & Parker was implemented in MATLAB (MathWorks, Inc., Natick, MA, USA) to estimate the risk level of the two dive exposures and is described in detail elsewhere.(20) To calculate the pDCS value for the two different exposures, the depth-time profile, breathing gas mix, and average weight for the study group were input to the Lillo & Parker model.

### Ultrasound Measurements

One week prior to the dive, each animal was anesthetized with isoflurane and the thorax was depilated via shaving cream and razor. Apical four-chamber cardiac (15–29 MHz, Vevo F2 system, FUJIFILM VisualSonics Inc., UHF29x probe, 221 FPS) and cross-sectional kidney (30 MHz, Vevo2100, FUJIFILM VisualSonics Inc., MS400 probe, 11 FPS) B-mode ultrasound recordings were collected. Imaging gel was generously applied to the region, and the probe was gently positioned above the ribcage for the cardiac view. The kidney was identified by palpation, and the ultrasound probe was placed over the abdomen. For both views, the probe position was adjusted until the maximum cross section of the heart or kidney was identified. Effort was made to obtain identical imaging planes at each timepoint and the imaged kidney was consistent for all measurements per animal. Ultrasound recordings were collected one week pre-dive to minimize effects of anesthetics on dive physiology, and every 10 minutes for the duration of the post-dive observation period (90 minutes or less).

### Image Review and Analysis

Cardiac recordings were exported from the Vevo F2 system in audio video interleave (avi) format and reviewed by a grader skilled at evaluating VGE in human echocardiography scans. Videos were evaluated at half speed playback or slower for identification of VGE presence. The Eftedal-Brubakk (EB) grading scheme(21) was adapted for rodents and used to assign an ordinal grade (0–5) based on VGE presence over 10 consecutive cardiac cycles. The scale was adapted only in the grade four description, in which units of 2 mm^2^ were used in place of cm^2^ as a more anatomically relevant unit of measurement (complete definitions in Table 1). The peak EB grade per animal was identified to characterize the presence of VGE in the heart throughout the post-dive observation period.

**Table 1:** The ordinal grading scale used for rodent echocardiogram assessment post dive, developed for this study, adapted from (21).

| *Grade* | *Semantic* |
| --- | --- |
| *0* | *No bubbles visible* |
| *1* | *Occasional bubbles* |
| *2* | *At least 1 bubble / 4 cardiac cycles* |
| 3 | *At least 1 bubble every cardiac cycle* |
| 4 | *At least one bubble at every 2mm<sup>2</sup> in every image* |
| 5 | *Whiteout, no single bubble discrimination* |

Ultrasound recordings of the kidney were offloaded in avi format and visualized for potential bubble presence and positional consistency. The total collected frames were trimmed for motion artifacts. In MATLAB, a region-of-interest (ROI) was manually drawn around the kidney and an average of the ROI pixel values was obtained for the remaining frames at each timepoint.

### Data Analysis

Model-estimated probabilities of DCS (pDCS) were reported for each dive profile to characterize differences in predicted decompression risk. Peak EB grades were identified for each animal across post-dive measurements and summarized by breathing gas. Renal image brightness was grouped by concurrent cardiac EB grade to explore trends in echogenicity associated with VGE burden. No inferential statistical testing was performed given the exploratory nature and small sample size of this pilot study.

### Renal Histology

A separate set of four Sprague Dawley rats (2 male, 2 female) were obtained from Charles River Labs for experimentation. In this separate set of dived animals (n = 4, aged 9 - 12 weeks, average weight: 179.4±45g), non-imaged kidneys were harvested postmortem and examined histologically for evidence of intrarenal gas. The dive profile differed from the previous section, instead consisting of an awake exposure for 90 minutes at 583 kPa while breathing air, with a 5 minute compression time and 10 minute decompression. Following post-dive monitoring and euthanasia, kidneys were harvested, fixed in 10% formalin for at least 48 h, routinely processed to paraffin, sectioned and stained using hematoxylin and eosin (H&E) by the UNC Pathology Services Core Facility. Sections were qualitatively evaluated by an experienced veterinary pathologist (RSS). Given the small sample size, these analyses were intended to provide preliminary observations rather than a definitive histological assessment of post-dive renal pathology.

## Results

### Decompression Exposure

In total, eight exposures were completed with four rats to each of the two dive protocols. All animals tolerated the hyperbaric exposure and decompression phase, then were anesthetized within minutes of exiting the chamber. The Lillo & Parker model produced a pDCS of 13% and 0.1% for the air and heliox dives, respectively (Fig. 2). One of eight animals completed the full 90-minute observation period, while three reached early humane endpoints due to labored breathing. The remaining four animals exhibited spontaneous respiratory arrest during observation and were immediately euthanized according to protocol. In two animals on the air exposure, skin lesions that resembled blisters were identified on the thorax and appeared to increase in size and quantity during the observation period. The mean time to study termination was 70.25±13 minutes for air dives and 56.8±8 minutes for heliox.

**Figure 2.**
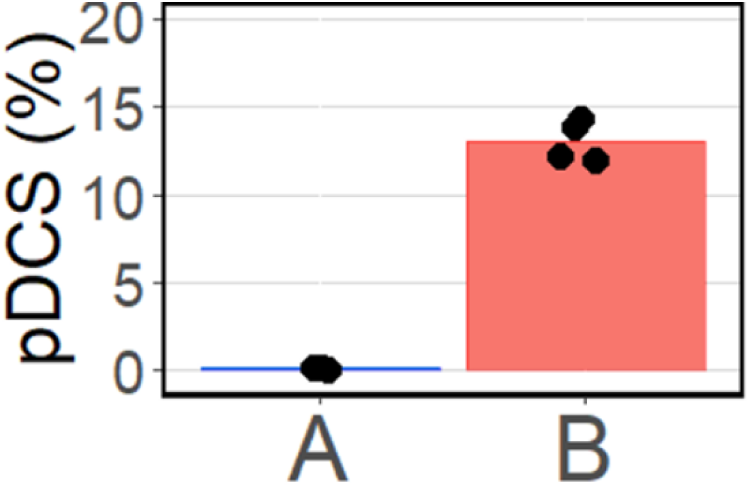
Probability of DCS (pDCS) calculated by the Lillo & Parker model(20) for rodents.

### VGE Detection

During cardiac imaging, multiple instances of flashes were identified in real time within the venous chambers by the researchers (Fig. 3a). These occasions were confirmed to be VGE during offline review, and additional instances were identified. VGE presence was variable, ranging from EB0 to EB4. The air dive was found to have a median peak grade EB2 (range: 1-4) and the heliox dives had a median peak grade EB1 (range: 0-1) (Fig 4).

**Figure 3.**
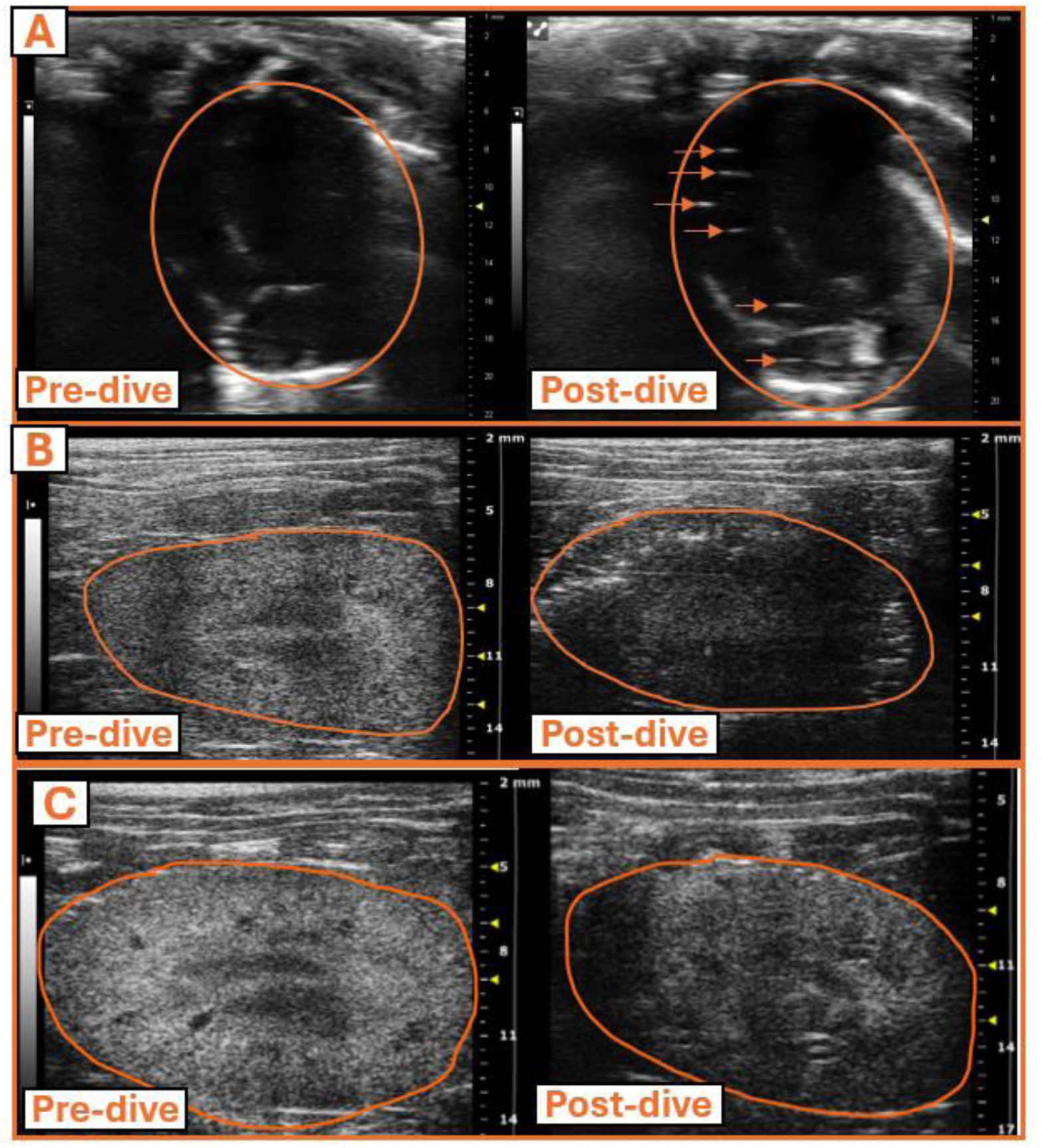
Pre- and post-exposure images of rats that completed an air dive. Panel (a) highlights the presence of venous gas emboli (VGE) in the venous cardiac chambers and (b) and (c) represents the images of the kidney with shadowing post-dive.

**Figure 4.**
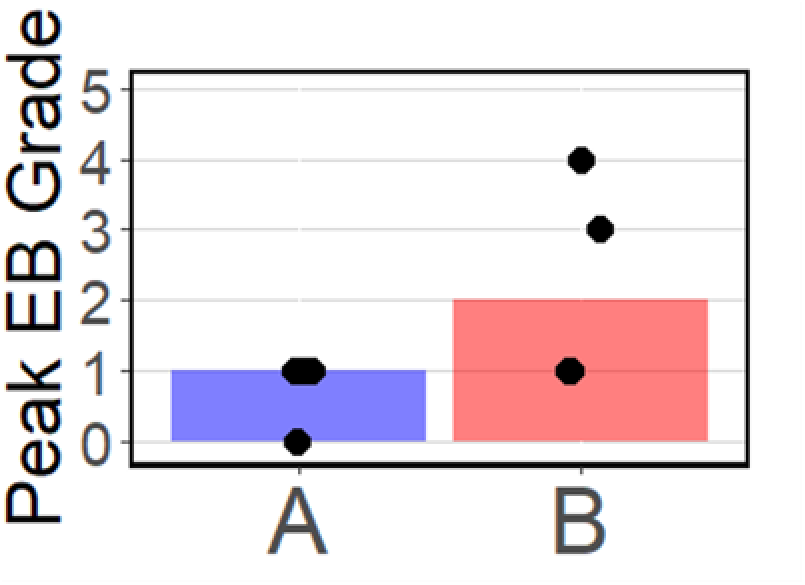
Peak EB grade identified per animal, separated by dive exposure (A represents Heliox, B denotes Air).

### Kidney Image Analysis

One clear instance of circulating gas bubble presence in the kidney was identified on the air profile, which occurred immediately prior to spontaneous respiratory arrest in the affected rat. Figure 3b shows the pre- and post- dive renal ultrasound of this rat, in which kidney ultrasound shadowing was noted. See figure 3c for a second example of shadowing in a separate rat. Comparison of kidney brightness and concurrent VGE cardiac presence revealed the timepoints with greater VGE presence (EB3-4) had a tendency for darker kidney images (Fig. 5).

**Figure 5.**
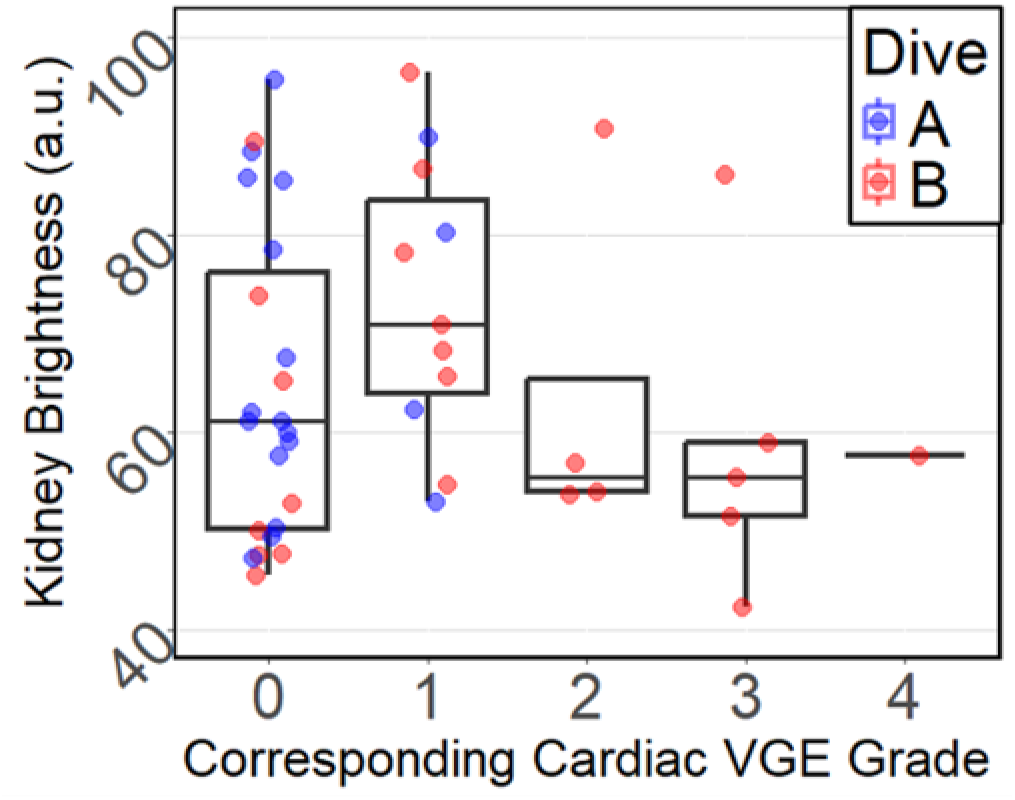
Concurrent evaluation of kidney image brightness and cardiac EB grade. Gas mixed denoted by A: Heliox and B: Air.

### Renal Histology

As a preliminary exploratory analysis, renal histology was performed in a separate cohort of four dived animals. All animals completed the full 90-minute post-dive observation period without reaching predefined humane endpoint criteria and were subsequently euthanized according to protocol. The Lillo & Parker model produced a pDCS of 0.2% for the dive completed by this subset of four animals. Because of the limited sample size, findings are reported descriptively. One animal of the four developed cutaneous vesicles and was found to have round voids in multiple arterioles of both kidneys (interpreted to be pre-mortem gas bubbles) and red blood cells in the tubules (interpreted to be secondary to capillary injury) (Fig. 6). No other abnormalities were identified in the other 3 kidney samples.

**Figure 6.**
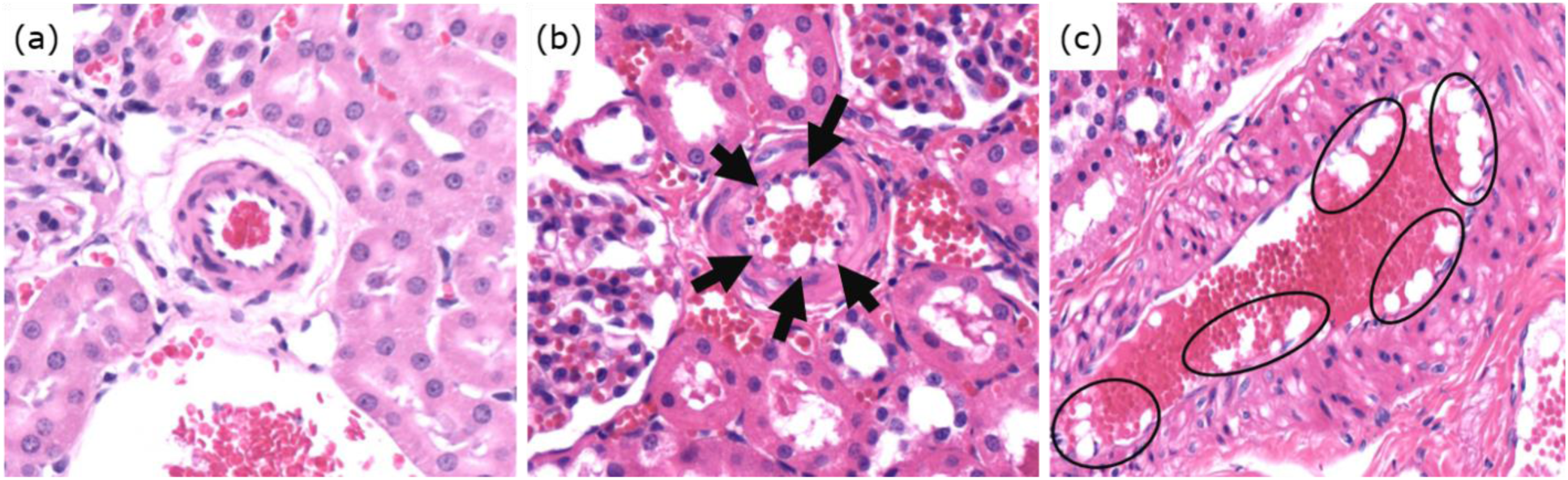
Post-dive rat renal histology samples stained using hematoxylin and eosin. (a) Normal arteriole containing red blood cells. (b) and (c) Arteriole containing clear round spaces within the lumen of the vessels along the endothelial surface. Clear round intravascular spaces interpreted as premortem gas bubbles are highlighted by arrows or outlined by black ovals. Both (b) and (c) images are from the same animal, which had skin lesions post-dive.

## Discussion

The decompression research field has focused on VGE in the venous lumen, extrapolating to the plethora of symptoms and regions involved in DCS pathophysiology.(1) However, it is theorized that bubbles form at the tissue level prior to circulating in the vasculature.(1,22) Further, the prevalence of VGE after dive exposures without DCS symptoms indicates that VGE typically do not cause injury.(23–25) An ability to detect and monitor bubble presence throughout the body could provide a basis to improve inert gas kinetic modeling. We have demonstrated the first concurrent multi-organ assessment in rodents looking at both cardiac VGE and renal ROI. The multi-point approach allowed for comparison against the standard human measurement (VGE) and correlation with a pDCS model. We found a decrease in kidney echogenicity over the post-dive monitoring period that inversely trended with an increase in concurrent cardiac VGE grade. We theorize that image shadowing may have occurred in the darkened kidney images, which we attribute to large gas presence superficial to or inside the kidney region. Previous preclinical studies by Currens *et al.* and Eltz *et al.* have found similar decreases in region brightness (acoustic shadowing) that were associated with severe DCS and high gas burden in swine and turtles, respectively.(17,18) Conversely, L’Abbate *et al.* identified an increased brightness in the liver of rats after hyperbaric exposure, which was attributed to gas bubble presence.(15) Expanded data collection is therefore necessary to confirm the cause of darkened kidney images; however, an ability to estimate tissue bubble presence could greatly change decompression stress estimation and move the field towards personalized monitoring.

Previous publications that report rodent ultrasound detection of VGE do not include multiple organ assessment.(5) In a recent scoping review, just one of 47 rodent decompression studies with biomedical imaging included a representative ultrasound image of their target organ with VGE.(5,26) The lone example ultrasound image from Rasdal *et al.* included a still frame of the pulmonary artery with bright spots, representing VGE.(26) In the present study, we documented VGE as similar bright spots in the venous chambers in apical four-chamber view and renal VGE from the transverse view. The air dives were found to have a higher pDCS level and greater VGE presence than the heliox dives, corresponding with previous DCS outcome in an awake rodent study group.(27) Orsat *et al.* exposed rats to the same dive profiles as the present study, but animals were kept awake for post dive DCS assessment, although without ultrasound imaging. In their study, the air dive resulted in 10 of 12 animals having severe decompression sickness, while four of 12 on the heliox profile had severe DCS. Inhaled anesthetics likely masked gait-related DCS symptoms in the current study, making it difficult to compare our animal outcomes directly to the Orsat *et al.* study.(27) However, our median peak VGE grade was higher in air than heliox dives, trending positively with the pDCS values and the severe DCS trends reported by Orsat *et al*.(27) This suggests that peak VGE measurements in rats may be risk sensitive at the population level, similar to humans.(4,28) The improved safety of heliox dives for rodents supports extensive prior work by Lillo, finding heliox reduced mortality.(29)

In this pilot study, only one rat completed the 90-min observation period without reaching an early endpoint; the remaining seven exhibited labored breathing or spontaneous respiratory arrest. Evaluating accuracy of pDCS from the Lillo & Parker model was not the primary objective, but was in accordance with a previous review finding the pDCS model to underestimate risk.(5) Additionally, the current study design was not powered to measure DCS rates across the study groups, limiting the direct connection between VGE measurements and DCS. Anesthesia may have influenced the onset and severity of adverse outcomes in both groups, and likely affected the time to study termination values for individual animals. However, anesthesia was a necessary tradeoff to ensure immobilization for consistent probe positioning and reliable longitudinal ultrasound measurements, as the primary objective was to develop imaging biomarkers of decompression stress rather than assess DCS outcomes.

An unexpected opportunity from depilation for ultrasound imaging was the identification of skin lesions during some post-dive observations. When present, skin lesions were identifiable in shaved areas and resembled blisters, which we theorize may be due to transcutaneous gas diffusion. Previous rodent DCS trials report qualitative symptom identification, but skin manifestations are rarely identified. It is possible that skin lesions have been under detected in rodent trials due to the dense fur. In the present study, skin lesions and high VGE presence were found to coincide, potentially indicating that both can occur under excess decompression stress. In humans, skin manifestations associated with DCS include mild rash, urticaria, and cutis marmorata.(30,31) Garcia *et al.* investigated four cases of cutis marmorata using ultrasound and found bubbles in the vasculature of the afflicted skin in all cases.(31) Additionally, all four patients in that study presented with other DCS symptoms and received recompression treatment, after which the lesions subsided.(31) Also in humans, Blenkarn *et al.* found skin lesions apparent after a switch in breathing gas at depth, likely due to local gas concentration differentials within the skin.(30) Taken in the context of the literature, our observation of skin lesions coinciding with higher VGE are relevant in the discussion of systemic bubble dynamics in this rodent model.

Separately, we identified an instance of intravascular gas bubbles in the histologically preserved kidney of a dived rat. Gas bubbles formed pre-mortem were highlighted upon veterinary review of the slides. Previous studies have found gas emboli in gross analysis of tissues post-dive, as well as in histological results. In a rodent model, L’Abbate *et al.* found that in animals that spontaneously died following a rapid decompression, bloody foam was visible upon dissection of the liver and gas emboli were present in histological liver samples.(32) In animals that reached humane endpoints later after decompression, fewer bubbles were observed both grossly and histologically.(32) In bycaught sea turtles, García-Párraga *et al.* found gas emboli present grossly in cardiac, stomach, and intestinal dissection, as well as in histological images of cardiac, kidney, and lung tissue postmortem in animals that had been diagnosed with moderate to severe gas emboli via radiographic and ultrasound imaging.(33) These previous findings emphasize the role of gas presence in the DCS mechanism, as well as how clinical diagnoses from medical imaging or observation can be corroborated with postmortem dissection.

A further limitation of this pilot study was the sample size. The 8 animal-dives performed as part of this pilot study necessitate further testing for definitive relationships to be established between VGE presence, kidney gas presence or other organ analysis, and DCS outcome. Furthermore, the described histological result was not a comprehensive finding, as one animal was found with evidence of tissue gas presence while three others did not. This result provides evidence that gas bubbles in organs can be visualized postmortem, and future work should include similar analysis. Further analysis to this end could corroborate acoustic findings (bright spots or shadowing) with microscopic histological findings.

## Conclusions

The developments in this study will serve as a foundation for future preclinical decompression research. This platform can be expanded for translational testing of advanced ultrasound techniques, while also collecting the standard VGE measurement. In this work, we found the dive with a higher pDCS from the model also resulted in a higher median VGE grade. Further, animals with high VGE presence tended to have darker kidney images, possibly attributable to acoustic shadowing from large bubble presence in the tissues. Separately, an instance of gas emboli was identified through histopathology, supporting the notion of acoustic shadowing. Future experimentation will include additional imaging locations to refine multi-organ assessment and tissue bubble detection, a necessary step towards physiologically relevant decompression models and wearable ultrasound for divers.

